# A role for Gcn5 in heterochromatin structure, gene silencing and NER at the *HML* locus in budding yeast

**DOI:** 10.1101/2021.02.28.433071

**Authors:** Hua Chen, Ling Zhang, Qikai Wang, Chenxi He, Lauren Frances Dender, Feng Gong

**Author notes:** To whom correspondence should be addressed: Dr. Feng Gong, Department of Biochemistry and Molecular Biology, University of Miami Miller School of Medicine, Miami, Florida 33136, Dr. Hua Chen, Department of Biochemistry and Molecular Biology, University of Miami Miller School of Medicine, Miami, Florida 33136.

## Abstract

Histone acetyltransferase Gcn5 plays an important role in transcription activation, DNA replication-coupled nucleosome assembly and nucleotide excision repair (NER). However, its functions on the heterochromatin are unexplored. Here, we find that removal of Gcn5 leads to more condensed heterochromatin structure, as revealed by topology analysis of *HML* circles. Importantly, the altered heterochromatin structure is restored by re-expression of Gcn5 in the *gcn5Δ* cells. As a result of the more compact heterochromatin, gene silencing at the *HML* locus is increased and NER efficiency at *HML* is impaired in the absence of Gcn5. Interestingly, while the association of SIR complex with *HML* is enhanced in cells lacking Gcn5, the altered compaction of *HML* heterochromatin is also observed due to the deletion of Gcn5 from Sir^−^ cells. These findings reveal a role of Gcn5 in the regulation of heterochromatin structure, gene silencing and NER efficiency at the heterochromatic *HML* locus in yeast.

## Introduction

Genome in eukaryotes is compacted to chromatin. The degree of chromatin compaction is not homogenous across the genome, leading to the existence of highly condensed regions known as heterochromatin or silenced chromatin and less condensed regions referred to as euchromatin (1,2). While gene expression is largely inactive, heterochromatin plays important roles in gene regulation and genome stability (3). Alerted heterochromatic state can impair gene expression patterns, leading to the development of different human diseases (4). The transcriptionally silent *HM* loci including *HML* and *HMR* located on chromosome Ш are a well-known heterochromatin locus in budding yeast *Saccharomyces cerevisiae*. Formation and maintenance of silenced *HM* loci require the silent information regulator (SIR) complex composed of Sir2, Sir3 and Sir4 (5). In the absence of SIR complex, *HML* heterochromatin is fully derepressed (6). On the other hand, overexpression of SIR complex is toxic to yeast cells by causing chromosome loss (7). In addition to the SIR complex, several other proteins have been identified to be involved in formation and/or maintenance of heterochromatin structure of *HML*, such as the chromatin remodeler Fun30 (6) and the NER (Nucleotide Excision Repair) protein Rad4 (8).

Gcn5 (General Control Non-depressible 5) is one of the best characterized histone acetyltransferase (HAT), belonging to the GNAT (Gcn5-related *N*-acetyltransferases) family of acetyltransferase enzymes (9). It functions in context of the transcriptional activator complex SAGA (10). Gcn5 acetylates histones at multiple positions including H3K9 and H3K14 (11–13), while recent studies indicated it can also acetylate many non-histone substrates (14,15). Moreover, acetyltransferase activity of Gcn5 to acetylate histones is weak on its own and the optimal activity requires Ada2 and Ada3 (16,17), another two subunits of SAGA complex. Ada2 physically interacts with Gcn5 and promotes its activity by enhancing binding of the enzymatic cosubstrate acetyl-CoA (18).

Gcn5 plays important roles in replication-coupled nucleosome assembly and DNA repair (19,20). It is generally considered to function on euchromatin because of the nature of being an acetyltransferase that activates transcription by acetylating nucleosomal histones (21). Currently its roles on heterochromatin are unclear. However, it has been reported that Ada2, the coactivator of Gcn5, is involved in transcriptional silencing at telomeres and ribosomal DNA (22). Furthermore, Gcn5-mediated histone acetylation is critical to efficient NER at a repressed yeast locus (11). These studies suggest a correlation between Gcn5 and the repressed or silenced chromatin. Here, we investigated the function of Gcn5 on *HML* heterochromatin structure. It was found that Gcn5 is involved in the compaction of *HML*. Removal of Gcn5 resulted in more condensed *HML* as revealed by topological analysis of *HML* circles. In addition, the heterochromatin in *gcn5Δ* cells is further silenced and NER of UV damage is impaired. These data identified Gcn5 as one more factor that contributes to the heterochromatin structure of *HML* and established a novel role of Gcn5 in budding yeast.

## Results

### Altered heterochromatin structure at the *HML* locus in the *gcn5Δ* cells

Chromosome structure can be reflected by the topology of DNA. To examine the structure of heterochromatic *HML* locus, we analyzed the DNA topology of *HML* using a method established in previous work (23). In this method, two recombination target sites (*FRT*) for the site-specific recombinase Flp1 were inserted in direct orientation at positions flanking *HML* (Fig. 1A, YXB2) or bracketing the coding region of *HML* and excluding the silencers (Fig 1A, YXB4). Galactose induction of Flp1 from a *GAL10-FLP1* construct resident elsewhere on chromosome leads to recombination between the two *FRT* sites and subsequent excision of *HML* as non-replicating mini-chromosome circles which are negatively supercoiled (8,23). Topoisomers of these *HML* circles can be separated by chloroquine-agarose gel electrophoresis. In the presence of 30 µg/ml chloroquine, the more negatively supercoiled circles migrate more slowly in the gel which can be visualized by Southern blotting (23). The topology of *HML* between different strains can be compared using the difference of linking number (ΔLk).

**Figure 1.**
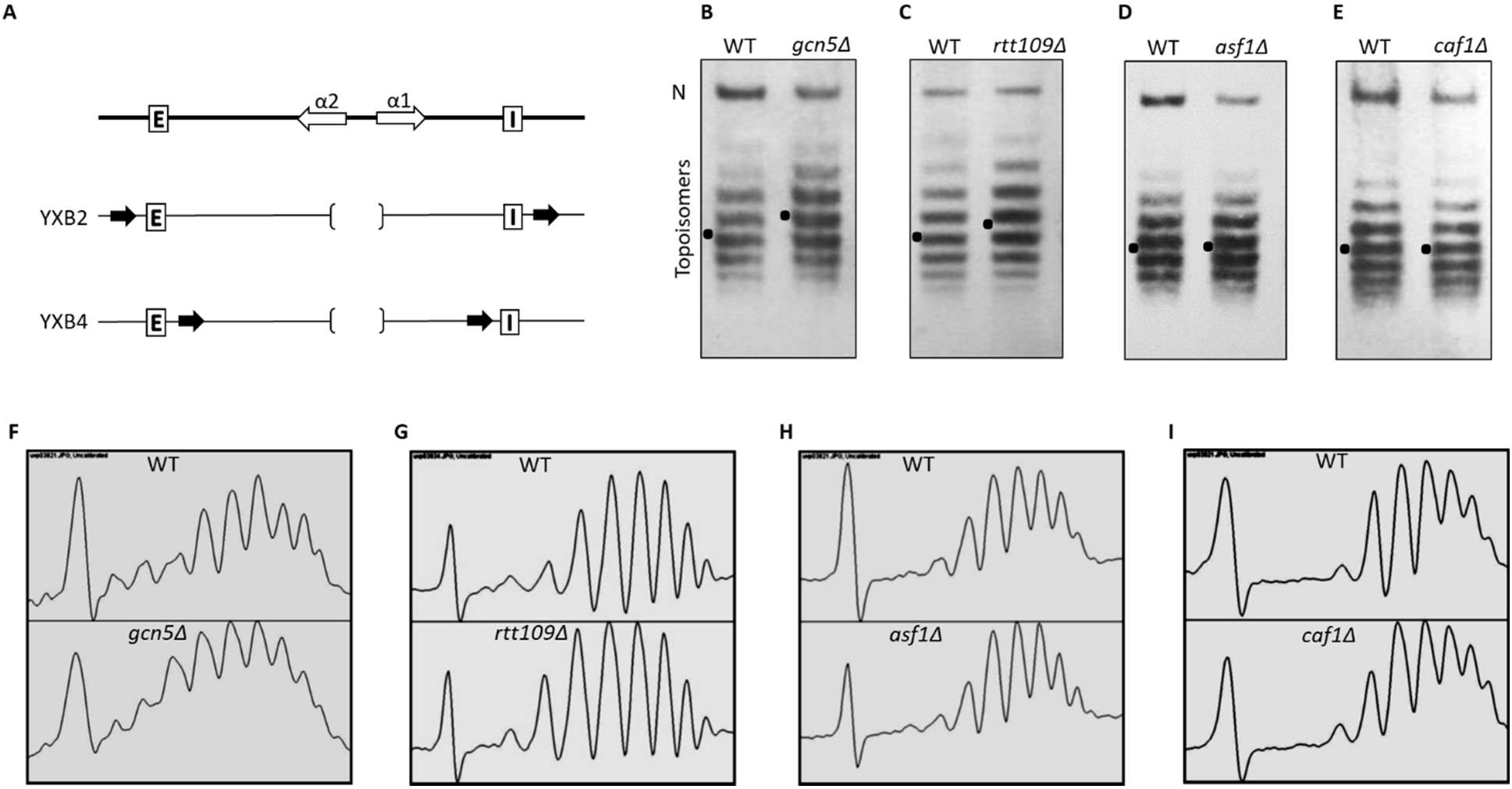
Analysis of *HML* circle topology in cells lacking Gcn5, Rtt109, Asf1 and Caf1. (A) Schematic representation of *HML* locus. *HML* in the original yeast strain (top) and in the two artificial yeast strains YXB2 and YXB4 (bottom) are shown. White open arrows, *α1* and *α2* genes; open squares, *HML* silencers E and I; black open arrows, *FRT* (FLP1 recombination target) sites. A deleted 294 bp fragment containing the divergent promoters of *α1* and *α2* is bracketed. (B-E) Topologies of *HML* circles isolated from the strains of YXB4 (WT), *gcn5Δ* (B), *rtt109Δ* (C), *asf1Δ* (D) and *caf1Δ* (E). A representative experiment carried out twice is shown. Cells of the indicated strains were grown to early log phase, followed by addition of galactose and additional incubation for 2.5 h. To detect *HML* circles, 40 µg of genomic DNA was fractioned by agarose gel electrophoresis in the presence of 30 µg/ml of chloroquine, subsequently transferred onto the membranes and subjected to Southern blotting as described in ‘Materials and Methods’. *HML* topoisomers, nicked circles (N) and the Gaussian center of topoisomer distribution (dots) are indicated. (F-I) Density profiles of the images B-E scanned by Image J software.

Inspired by our earlier work revealing that NER protein Rad4 regulates the heterochromatin conformation of *HML* and telomere (8), we set out to investigate whether other Rad proteins in the Rad3 epistasis group (24) including Rad1, Rad7, Rad14, and Rad 23 are involved in the regulation of *HML* structure. Results of Southern blots (Fig. S1) indicated that removal of Rad1, Rad7 and Rad 14 didn’t affect the DNA topology of *HML* circles. In contrast, deletion of Rad23 led to more negatively supercoiled *HML* circles, similar to that observed in *Rad4Δ* cells (8). Given Rad23 interacts with Rad4 forming the Rad4-Rad23 heterodimer which is crucial for NER (25), we assume the Rad4-Rad23 heterodimer plays a role in heterochromatin structure at the *HML* locus.

Because it has been known that histone modifications (26–28) and histone chaperones (29–31) play a critical role in heterochromatin silencing in yeast, we next examined the involvement of two histone acetyltransferases (Gcn5, Rtt109), and two chromatin assembly factors (Asf1, Caf1) in the regulation of heterochromatin structure at *HML* locus. As shown in Fig. 1B and 1F, topologies of *HML* circles in YXB4 (WT) and the *gcn5Δ* strain were different with ΔLk of ~1. Moreover, those circles isolated from the *gcn5Δ* cells were more negatively supercoiled, suggesting *HML* was more compacted in the *gcn5Δ* cells than WT. On the other hand, less topology changes were observed when the *RTT109* acetyltransferase was disrupted (Fig.1, C and G). Additionally, topologies of *HML* circles were not altered by removal of ASF1 and CAF1 (Fig. 1, D, E, H and I), suggesting these two histone chaperones are not involved in building up the heterochromatin structure of *HML* although they have been proven to contribute to the silencing of *HML* (29–31). Subsequent studies focused on Gcn5 which contributed most to *HML* structure among these four tested factors.

### Re-expression of Gcn5 in the *gcn5Δ* cells restores *HML* heterochromatin structure

To test if *HML* heterochromatin structure in *gcn5Δ* cell can be restored by re-expression of Gcn5, a low copy *CEN* vector carrying the *GCN5* gene under the control of its native promoter was introduced into wild type (YXB4) and the *gcn5Δ* cells. Topology analyses by Southern blotting (Fig. 2, A and B) indicated that the existence of empty vector in WT cells didn’t affect the topologies of *HML* circles (Fig. 2A, lane 2). In addition, overexpression of *GCN5* in WT cells didn’t alert the *HML* topology either (Fig. 2A, lane 1), suggesting cellular Gcn5 is abundant for maintaining the heterochromatin structure of *HML*. Importantly, re-expression of *GCN5* in *gcn5Δ* cells led to similar supercoiling of *HML* circles to that in wild type (Fig. 2A, lanes 2 and 4), restoring the altered heterochromatin structure of *HML* resulted from deleting *GCN5* (Fig. 2A, lane 3).

**Figure 2.**
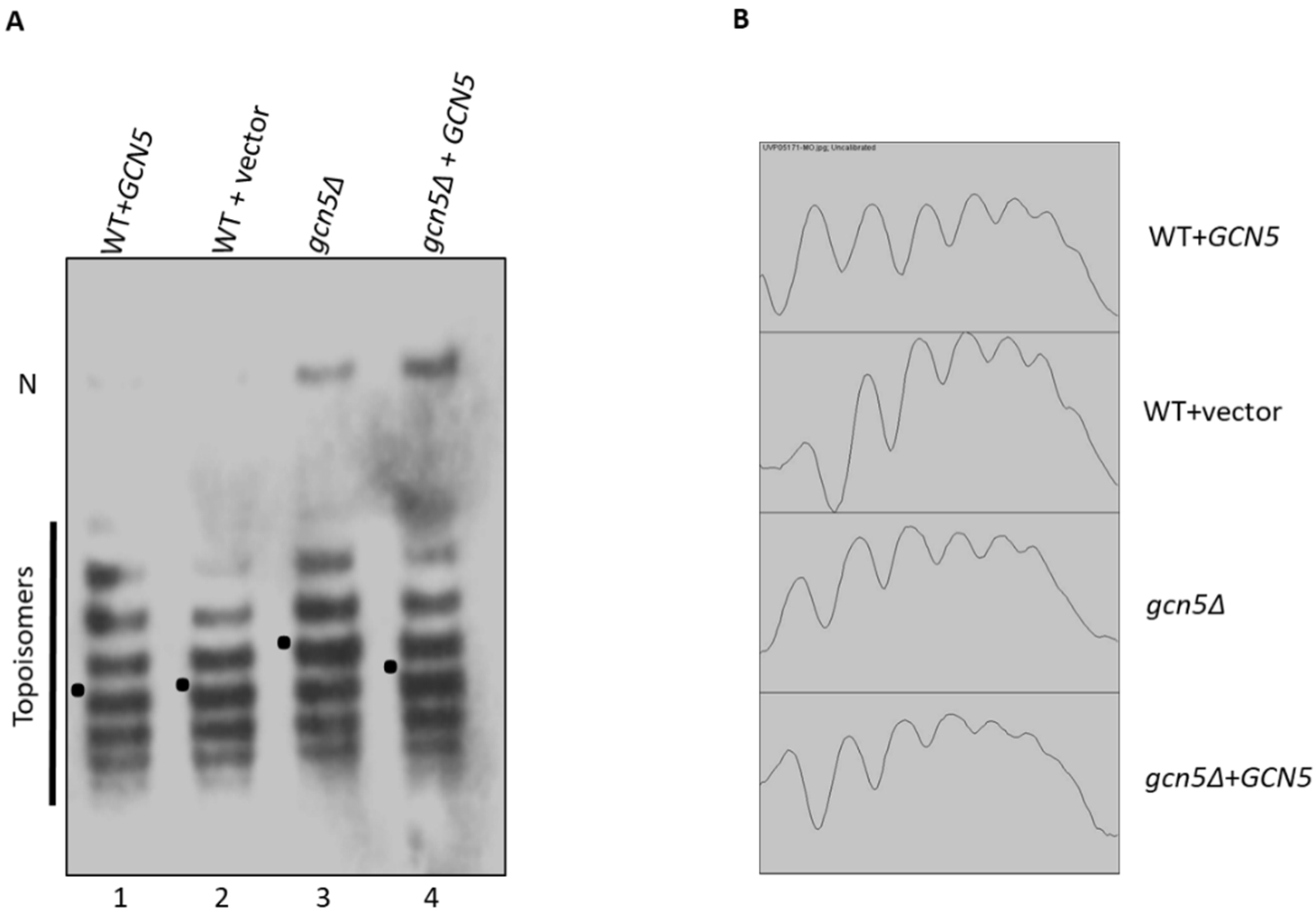
Topology analysis of *HML* circles in cells re-expressing Gcn5. (A) Topologies of *HML* circles detected by Southern blotting. Expression of *GCN5* was driven by its native promoter using a low copy expression vector. The empty vector or the vector taking *GCN5* gene was introduced into YXB4. *HML* circles were formed by the induced recombination between the two *FRT* sites. To compare *HML* topology of the indicated strains, 40 µg of genomic DNA isolated from the mid-log phase cells was subjected to Southern blotting as described in ‘Materials and Methods’. *HML* topoisomers, nicked circles (N) and the Gaussian center of topoisomer distribution (dots) are indicated. A representative experiment carried out twice is shown. (B) Density profiles of the image in A scanned by Image J software. The density profiles of topoisomers are presented excluding the nicked circles.

### Enhanced association of SIR protein to *HML* in the *gcn5Δ* cells

Since SIR complex is prerequisite for the establishment and the maintenance of *HML* heterochromatin structure, we next examined if deletion of *GCN5* altered the levels of *HML*-bound Sir2 protein. Equal amounts of whole cell extracts from YXB4 (WT), the *gcn5Δ* strain and the *sir3Δ* strain were subjected to ChIP assay using the specific antibody of Sir2. Subsequently the DNA fragments of four regions spanning from silencer E to silencer I on *HML* locus were analyzed by PCR. As shown in Fig. 3A, Sir2-bounded fragments were observed on all four detected regions of NucS, Nuc8, Nuc7 and Nuc3 in the *gcn5Δ* cells, indicating that removal of *GCN5* didn’t affect the spreading of SIR complex on *HML*. Furthermore, the amounts of all these four fragments were ~2.5 fold more in the *gcn5Δ* cells than in WT (Fig. 3A, B), while the results of Western blots indicated that the cellular amounts of Sir2 were slightly increased in the *gcn5Δ* cells (Fig. 3C). These data suggested that the levels Sir2 bound at *HML* were increased due to the deletion of *GCN5* from *SIR*^*+*^ cells. In addition, deletion of *SIR3* impaired the association of Sir2 at the regions of NucS, Nuc8 and Nuc7 (Fig. 3A).

**Figure 3.**
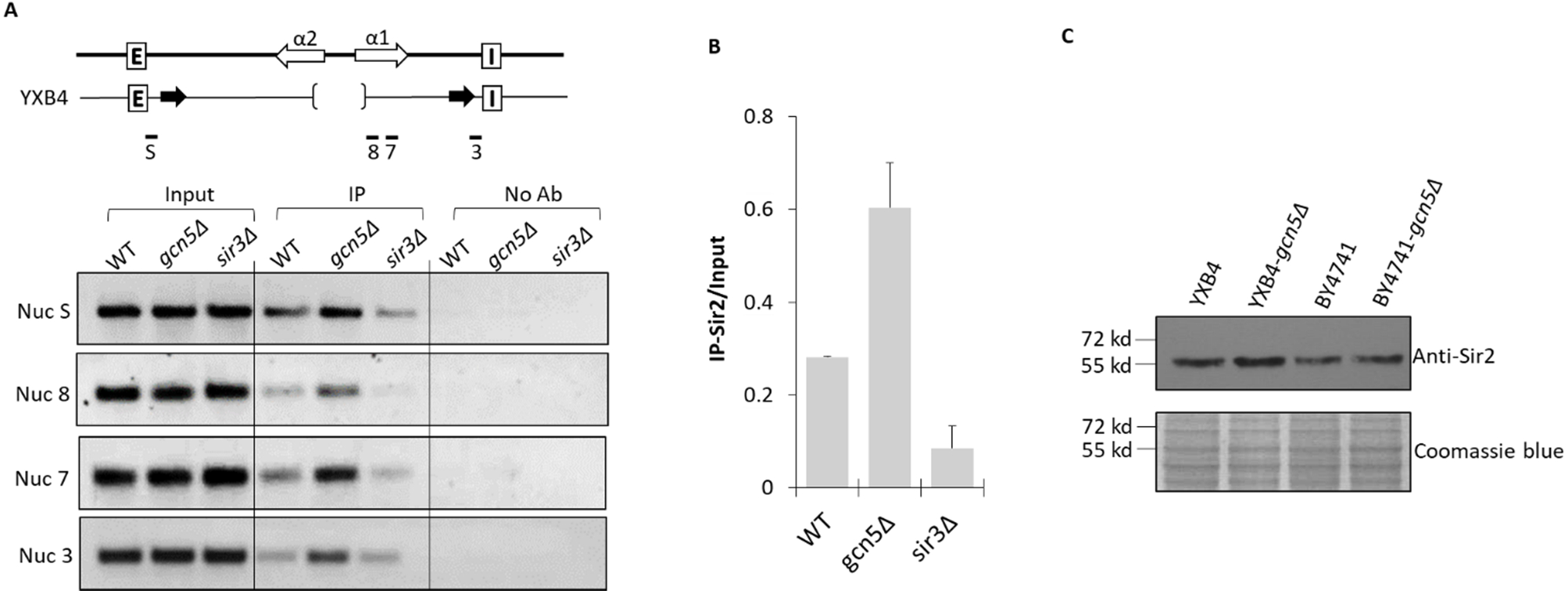
Examination of the levels of Sir2 bound at *HML* locus. (A) Top panel, schematic positions of the four tested regions on *HML* of YXB4; Bottom panel, association of Sir2 with *HML* detected by ChIP. The DNA fragments bound with Sir2 were isolated using Sir2-specific antibody, subsequently the amounts of the purified DNA fragments of four individual regions (NucS, Nuc8, Nuc7, Nuc3) were analyzed by PCR as described in ‘Materials and Methods’. A representative experiment carried out three times is shown. (B) Quantification of the data of Nuc7 in panel A. The levels of Sir2 bound at *HML* were determined by comparing the amounts of Sir2-associated DNA fragments (IP) with that of the total DNA fragments (input). The average ± S.D. from three independent experiments is presented. (C) Sir2 expression in total cell extracts. 15 µg of total protein from the two isogenic WTs (YXB4, BY4741) and the *gcn5Δ* strains was separated by 10% SDS-PAGE and subjected to Western blotting analysis. A representative experiment carried out three times is shown. Coomassie blue staining was used for the loading control.

### Adverse effects of Gcn5 and Sir3 on the *HML* circle topology

To further evaluate the coordination of Gcn5 and SIR complex on the regulation of *HML* structure, we compared the topology of *HML* circles isolated from the strains of *gcn5Δ, sir3Δ* and the double mutant *gcn5Δ sirΔ*. Because the *FRT* sequences were positioned differently in YXB4 and YXB2 (Fig. 1A), *HML* circles isolated from YXB4 (Fig. 4A, lanes 1-4) were smaller than those from YXB2 (Fig. 4A, lanes 5-8), which are 2.9 kb and 3.8 kb, respectively (23).

**Figure 4.**
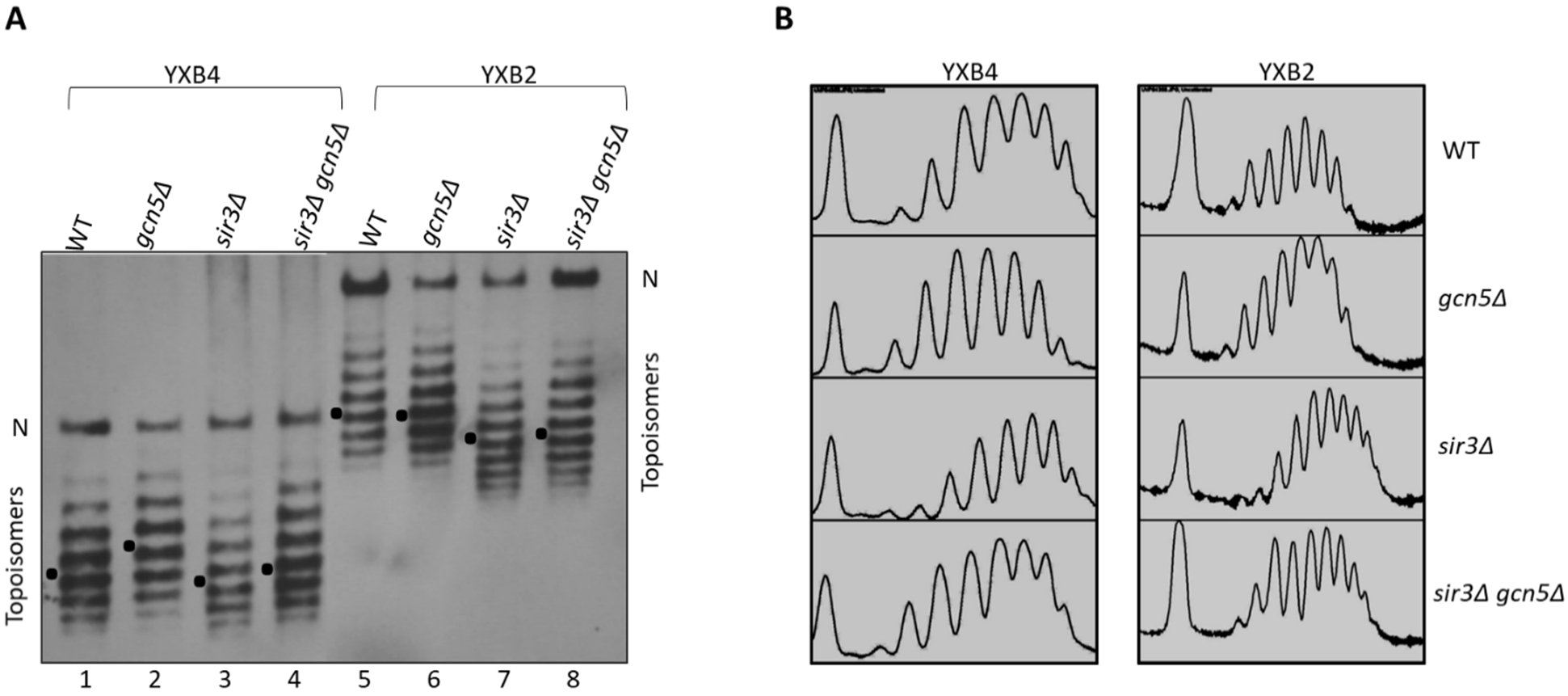
Comparison of the topology of *HML* circles in the *gcn5Δ* and the *gcn5Δ sirΔ* cells. (A) Detection of topology of *HML* circles by Southern blotting. 40 µg of genomic DNA from the indicated strains was fractioned in agarose gels supplemented with 30 µg/ml chloroquine and was subjected to Southern Blotting analysis. Two sets of genomic DNA samples isolated from the strains under two wild type (YXB4, YXB2) backgrounds were used. A representative experiment carried out twice is shown. *HML* topoisomers, nicked circles (N) and the Gaussian center of topoisomer distribution (dots) are indicated. (B) Density profiles of the image in A scanned by Image J software.

Deletion of *GCN5* from YXB4 led to more negatively supercoiled *HML* circles (Fig. 4A, lane 2), confirming the earlier observations (Fig. 1B and 1F). Adversely, deletion of *SIR3* resulted in total disruption of *HML* heterochromatin structure (Fig. 4A, lane 3), in agreement with previous work (23). Interestingly, the topology status of *HML* circles from the *gcn5Δ sirΔ* mutant was at an intermediate level between the status of the circles in the *gcn5Δ* and the *sir3Δ* cells (Fig 4A, lane 4). They were more condensed than the circles in *sir3Δ* cells and were less condensed than the circles in *gcn5Δ* cells (Fig. 4B, right panel). Therefore, removal of *GCN5* led to more compacted *HML* heterochromatin structure in both *SIR*^*+*^ cells and *SIR*^*-*^ cells of YXB4.

Consistently, the *HML* heterochromatin circles became relaxed without Sir3 in YXB2 cells (Fig. 4A, lane 7). However, deletion of *GCN5* from *SIR*^*+*^ cells and *SIR*^*-*^ cells of YXB2 didn’t lead to detectable alterations of *HML* (Fig. 4A, lanes 6 and 8). Since the two recombination target sites for the Flp1 recombinase were inserted in YXB2 at positions flanking *HML* and including the E and I Silencers, instead of bracketing *HML* and excluding the silencers in YXB4 (Fig 1A), a reasonable explanation is that released *HML circles with the silencers* of YXB2 diminishes or interferes with the function of GCN5 on the heterochromatin structure through an unknown mechanism.

### Transcriptional silencing was promoted in the *gcn5Δ* cells

Because the removal of *GCN5* let to a more compact heterochromatin structure and increased binding of SIR complex at *HML* locus, we assume that gene silencing at *HML* will be promoted in the *gcn5Δ* cells. To test this possibility, expression of the *URA3* reporter gene replacing *HMLα* mating genes of *α1* and *α2* (Fig. 5A) (32) were determined by measuring cell survival rate in medium containing 0.1% of 5-fluroorotic acid (FOA). As observed in previous work (33) and as shown in Fig. 5A, the *gcn5Δ* cells grew much slower than wild type (YXB61-1) in synthetic complete (SC) medium. Insertion of *URA3* at *HML* increased the distance between Silencers E and I, consequently *URA3* is not completely silenced and ~18% cells of wild type grew on FOA plates (Fig. 5A and 5B), in agreement with our earlier work (8). Importantly, the amounts of survived cells was increased to about 25% (~1.4 fold) in the absence of *GCN5*, indicating that the *URA3* gene was further silenced and the transcription of *URA3* was reduced in the *gcn5Δ* cells. These data suggested that deletion of *GCN5* enhanced gene silencing at *HML* locus.

**Figure 5.**
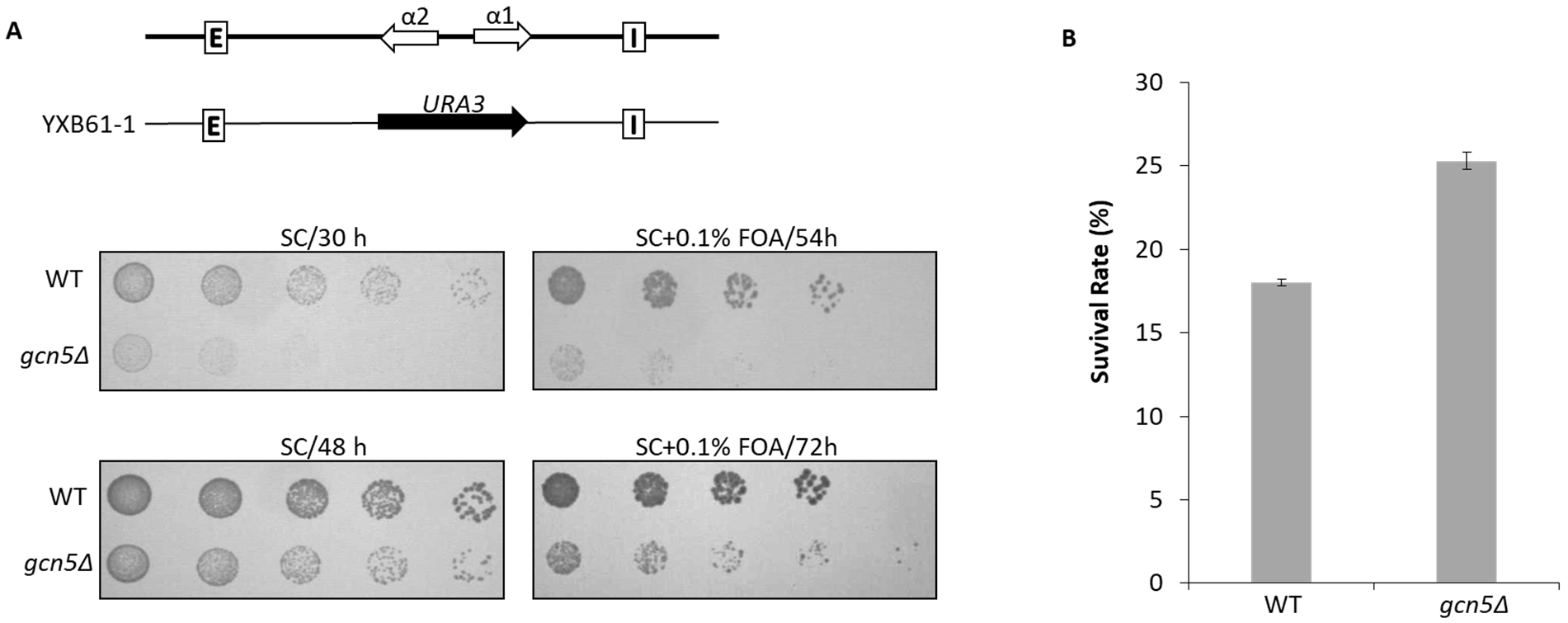
Examination of gene silencing at *HML* locus in absence of Gcn5. (A) Spot test assays of WT (YXB61-1) and the *gcn5Δ* cells. To test the gene silencing, a previous constructed strain YXB61-1 was used, in which the *HMLα* gene was replaced by the *URA3* gene (top panel). Expression of the *URA3* gene was monitored by spot test on SC plates and SC+0.1% FOA plates as described in ‘Materials and Methods’. A 4-fold serial dilution of the cultures of WT and the *gcn5Δ* strain was made, 10 µl of the diluted cultures was subsequently spotted out on the plates and incubated at 30°C for indicated time. A representative experiment carried out three times is shown. (B) Survival rate of WT and the *gcn5Δ* cells on the SC+0.1% FOA plates. To measure the survival rate of the cells on the FOA plates, the diluted cultures were spread on the plates and incubated at 30°C for 72 h. Colony numbers were counted using Alphaimager 2000 software. Survival rate was calculated as described in ‘Materials and Methods’. The average ± S.D. from three independent experiments is presented.

### Reduced NER efficiency in the *gcn5Δ* cells

Gcn5 has been reported to play an important role in NER at a few transcribed genomic regions or non-transcribed genomic regions (11,34–36), its role in NER in heterochromatin, such as the *HML* locus has not been investigated. To examine whether NER efficiency at the heterochromatic *HML* is altered in the *gcn5Δ* cells, YXB4 cells were irradiated with 200 J/m^2^ UV, followed by additional growth in the dark for NER. Efficiency of NER was monitored using a PCR-based method developed previously (37). Equal amounts of genomic DNA were used for the individual PCR reactions, as indicated by the equal amounts of PCR products from amplifying a short 0.3 kb *HML* fragment (Fig. 6A). When a 2.9 kb fragment was amplified, PCR efficiency reflected the NER efficiency because the existence of more DNA lesions on a longer DNA fragment blocks the Taq DNA polymerase. As a result, PCR products increased with the repairing time (Fig. 6A) and ~65% of the 2.9 kb fragment was repaired 2 h after UV treatment (Fig. 6B and 6C). In contrast, deletion of *GCN5* reduced the NER rate and ~45% of the *HML* region was repaired 2h after UV treatment (Fig 6B), 1.4-fold less than the wild type. These data indicated that Gcn5 is involved in the NER process at the *HML* locus. Additionally, removal of *SIR3* increased NER efficiency on the same 2.9 kb region of *HML* locus (Fig. 6C), consitent with a previous observation that NER efficiency at the repressed subtelomere region was enhanced in absence of *SIR2* (38).

**Figure 6.**
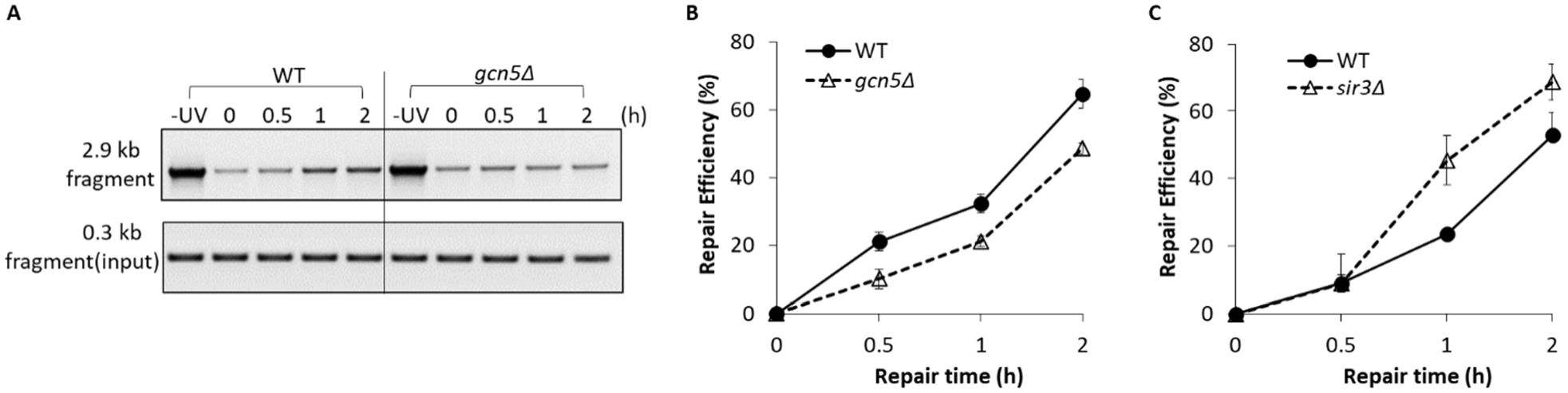
Evaluation of NER efficiency at *HML* locus in the cells lacking Gcn5. (A) Analysis of NER on a 2.9 kb fragment of *HML* in WT and the *gcn5Δ* cells. Cultures were harvested at the indicated time after 200 J/m^2^ of UV treatment, followed by isolation of genomic DNA and PCR analysis as described in ‘Material and Methods’. A 0.3 kb fragment of *HML* was amplified to monitor equal amounts of genomic DNA was used. (B) NER efficiency on the 2.9 kb fragment of *HML* in the *gcn5Δ* cells. The data in panel A was quantified and the NER efficiency was calculated as described in ‘Materials and Methods’. (C) NER efficiency on the 2.9 kb fragment of *HML* in the *sir3Δ* cells. For both B and C, the average ± S.D. from three independent experiments is presented.

## Discussion

The data presented here revealed a role of the histone acetyltransferase Gcn5 in the regulation of heterochromatin structure, DNA repair and gene silencing at the *HML* locus. We found that removal of Gcn5 led to more compact heterochromatic *HML* and enhanced gene silencing. Furthermore, we revealed that Gcn5 is involved in regulating the association of SIR complex with *HML* and in regulating NER efficiency of the heterochromatin. These findings shed light on a novel function of Gcn5 on heterochromatin beyond its role on euchromatin as a transcription activator.

It is likely that the functions of Gcn5 on the structure and the gene silencing of *HML* locus are related to its activity of acetylating nucleosomal histones. In SIR^+^ cells of YXB4, Gcn5 regulates the amounts of SIR bound with *HML* which is critical to the conformation of the heterochromatin (5,8). Given that Gcn5 is a histone acetyltransferase and that SIR complex bind preferentially to unacetylated histones (2), it is possible that removal of Gcn5 leads to further hypoacetylation of *HML* locus which increases the binding of SIR complex. Furthermore, the assumption suggests that there do exist certain levels of acetylated histones targeted by Gcn5 at *HML* locus to prevent heterochromatin hypercondensation, while most other histones at the heterochromatin are deacetylated by SIR complex (13,39). Therefore, it is reasonable that SIR complex doesn’t deacetylate all histones acetylated by Gcn5. Indeed, Gcn5 can acetylate histones at many different positions of histone H3 and H4 (13,16,40,41), and yet Sir2 specifically targets H3K9, H3K14 and H4K16 (42,43). Importantly, the viewpoint is directly strengthened by the observation that deletion of Gcn5 alters the heterochromatin structure of SIR^−^ cells of YXB4 (Fig. 4A). Taken together, we presume that Gcn5-mediated histone acetylation regulates the conformation and gene silencing at the heterochromatic *HML* locus.

Studies on NER at the repressed *MFA2* promoter in budding yeast provides a significant view of how Gcn5 is involved in the NER process. Following UV irradiation, GG-NER (Global Genomic NER) complex Rad7-Rad16-mediated increase of Gcn5 occupancy at *MFA2* promoter leads to elevated levels of histone acetylation of H3K9 and H3K14, which promotes an open chromatin structure at *MFA2* and consequently increases NER efficiency (11,20,44). Our primary observations indicate that only basal levels of Gcn5 occupancy was observed at *HML* prior to UV treatment, whereas the occupancy was enhanced dramatically after UV irradiation (data not shown). Notably, we found that deletion of Gcn5 impaired the NER efficiency at *HML* (Fig. 6B). However, the mechanism that Gcn5 contributes to NER at heterochromatic *HML* is likely different from that at *MFA2* promoter, since UV irradiation is unable to stimulate histone H4 or H3 acetylation at the repressed subtelomere due to the presence of Sir2 (38). Hence, how Gcn5 is involved in NER of *HML* locus needs to be further investigated.

In summary, Gcn5 plays an important role in the structure maintenance, gene silencing and NER process at the *HML* heterochromatin. These findings suggest that the function of Gcn5 at the heterochromatin correlates with its acetyltransferase activity targeting other histones instead of histone H3K9 and H3K14 at *HML* locus, which are deacetylated by the SIR complex. It is also possible that heterochromatin is more dynamic than we originally thought, there are always active competitions between the Gcn5 histone acetyltransferases and the SIR deacetylase at the *HML* heterochromatin. Thus, it will be of considerable interest to profile a global map of histone acetylation at the heterochromatic *HML* in order to further evaluate the contribution of Gcn5 to the heterochromatin.

## Materials and Methods

### Materials

Pierce™ high sensitivity streptavidin HRP and protein A agarose were purchased from Thermo Fisher Scientific™. Primary antibody of Sir2 was from Santa Cruz Biotechnology. Yeast synthetic complete medium were purchased from Clontech. Taq DNA polymerase was from Sigma Aldrich. Amersham hybond-N^+^ nylon membrane was obtained from GE Healthcare Life Sciences. Oligonucleotides used in this work were synthesized by Sigma Aldrich. All other chemicals are molecular biology grade.

### Yeast strains

Yeast strains of YXB2, YXB4, YXB2 *sir3Δ*, YXB4 *sir3Δ* (23) and YXB61-1 (32) were provided by Dr. Bi (University of Rochester). Single deletion mutants of Rad proteins and Gcn5 were from laboratory stock. The double mutants of YXB2 *gcn5Δsir3Δ* and YXB4 *gcn5Δsir3Δ* were constructed in this work by removing GCN5 from the *sir3Δ* mutant using PCR-based gene deletion strategy (45,46). All strains were confirmed by DNA sequencing. Absence of *GCN5* were complemented by re-expressing *GCN5* cloned into the expression vector YEp352 (47). *GCN5* gene was amplified from genomic DNA of wild type (YXB4) using the primers HC01 and HC02 (Table S1).

### Topological analysis

Topological structure of *HML* locus was analyzed as previously described (8). Yeast cells were grown to early log phase (OD600=0.6) in YPR medium (10 g/L yeast extract, 20 g/L peptone, 20g/L raffinose) at 30°C. Galactose (2%) was added to the culture that was further incubated for 2.5 h to induce the expression of Flp1 recombinase under control of *GAL10* promoter. Genomic DNA was isolated using the glass bead method and was fractioned on 1.2% agarose containing 30 µg/ml of chloroquine, followed by transferred to the positively charged nylon membranes. *HML* circles were detected by Southern blotting using a *HML*-specific probe (5’-biotin-TCTTCTTGAGACAATTTGGCC-3’) complementary to the *α1* gene coding region (8). The 5’-biotin labeled probe was incubated with membranes overnight at 45°C. Membranes were washed twice with buffer I (2 × SSC, 0.1% SDS) and buffer II (0.2 × SSC, 0.1% SDS) before blocked using 3% BSA prepared in PBST buffer. To detect *HML* circles, membranes were incubated with high sensitive streptavidin HRP overnight at 4°C, followed by detection with ECL reagents. The Gaussian center of topoisomer distributions was determined as described previously (8).

### Chromatin immunoprecipitation (ChIP)

ChIP assays were performed as described in earlier work (8). Mid-log phase yeast cells were treated with 1% formaldehyde for 15 min at room temperature, pelleted, and washed twice with TBS (25 mM Tris, pH 7.5, 137 mM NaCl, 2.7 mM KCl). Crosslinked cells were suspended in lysis buffer (50 mM HEPES–KOH, pH 7.5, 140 mM NaCl, 1 mM EDTA, 1% Triton X-100, 0.1% sodium deoxycholate, 1 mM PMSF, 1 mg/ml leupeptin, 1 mg/ml pepstatin A) and disrupted using glass beads, followed by sonication to yield DNA fragments with an average size of 300 bp. Protein concentration were measured by Bradford assay. Equal amounts of protein from each sample were incubated with specific antibody for Sir2 overnight at 4°C. The immunocomplex was precipitated using Protein A sepharose beads. The beads were consecutively washed with the lysis buffer, wash buffer 1 (lysis buffer containing 500 mM NaCl), wash buffer 2 (10 mM Tris-HCl, pH 8, 250 mM LiCl, 0.5% NP-40, 0.5% sodium deoxycholate, 1 mM EDTA) and TE buffer and then treated with RNase A in TE at 37°C for 30 min. Chromatin was then eluted from the beads using elution buffer (1% SDS, 0.1 M NaHCO3) and the cross-link reversed by incubation at 65°C overnight. DNA was then deproteinized by the addition of 4 μl of 10 mg/ml proteinase K and incubation at 37°C for 2 h. After phenol–chloroform extraction and ethanol precipitation, DNA was resuspended in 20 μl of TE and was used as the templates for PCR assays. Total four regions of NucS, Nuc8, Nuc7 and Nuc3 were analyzed using *HML* specific primers. Respectively, NucS was amplified using the primers HC03 and HC04; Nuc8 was amplified using the primers HC05 and HC06; Nuc7 was amplified using the primers HC07 and HC08; Nuc3 was amplified using the primers HC09 and HC10. Nucleotide sequences of all these primers were listed in Table S1. PCR products were resolved on 1.5% agarose gels.

### Western Blotting

Total proteins were extracted from mid-log phase cells of WT and *gcn5Δ*. Protein concentration was measured by Bradford assay. 15 µg of proteins were separated by 10% SDS-PAGE and were transferred to nitrocellulose membranes. Protein levels of Sir2 were detected using the protein-specific primary antibody.

### Analysis of NER efficiency on *HML* locus

Cells of wild type and the *gcn5Δ* strain were grown to early-log phase in YPD medium at 30°C. A part of cultures were harvested prior to UV treatment, the remained cultures were irradiated with 200 J/m^2^ UV light and were incubated additionally for different time (0 h, 0.5 h, 1 h and 2 h) in the dark. Genomic DNA isolated from UV-treatment samples and control samples were used as PCR templates for analyzing NER efficiency at *HML* locus using a PCR-based technique developed previously (37). Rational of this technique is that DNA lesions on templates are able to block movement of PCR polymerase, resulting in decreased amplification. To monitor the process of DNA damage repair, a 2.9 kb fragment of *HML* was amplified from each sample using the primers HC03 and HC10 (Table S1). A 0.3 kb fragment of *HML* was amplified using the primers HC03 and HC04 (Table S1) to confirm equal amounts of genomic DNA was added into each PCR reaction. PCR products were detected on 1.0-1.5% agarose gels and were quantified using Image J software. Efficiency of DNA damage repair at the 2.9 kb fragment was defined as the percentage of [(amounts of PCR products at each time points)/(amounts of PCR products of control)], in which the amounts of the 2.9 kb fragment were normalized using amounts of the 0.3 kb fragment.

### Analysis of gene silencing at *HML* locus

Yeast cells of YXB61-1 and YXB61-1 *gcn5Δ* strains were grown to mid-log phase in YPD medium. To perform the spot assay, the cultures were first diluted to OD600=0.005, followed by making a 4-fold serial dilutions. 10 µl of each diluted samples were spot out on the SC plates and the SC+ 0.1% FOA (5-fluoroorotic acid) plates. To calculate the cell survival rate, cultures were harvested by centrifuge and were resuspended in SC medium. The diluted cultures were spread on the SC plates and the SC+ 0.1% FOA plates and incubated at 30°C for 72 h. Colony numbers were counted using Alphaimager 2000 software. Survival rate was defined as the percentage of [colony numbers on SC+ FOA plates]/ (colony numbers on SC plates)].

## Acknowledgments

We thank Dr. Bi (University of Rochester) for providing us yeast strains. This work was supported by R01 ES017784 (to FG) from NIH.

