## Supplementary material for "A role for Gcn5 in heterochromatin structure, gene silencing and NER at the *HML* locus in budding yeast": Table S1, Fig. S1

Oligonucleotides used in this study.

| Oligo | Sequence (5'-3') |
|---|---|
| HC01 | AATGAGCTCCAAGGCAATTATTG |
| HC02 | AATGGTACCCTTCGAAAGGAATAGTAGC |
| HC03 | AGTTTTCGGCACGGACTTATTTGG |
| HC04 | TCGTCTAATACAAGTTTGAATGACG |
| HC05 | AATCATACAGAAACACAGC |
| HC06 | AAATCGAGAGGAAGGAAC |
| HC07 | CAGGATAGCGTCTGGAAGT |
| HC08 | GACATTTTGTTTTACACCG |
| HC09 | ACTTTCGTATTTAAGGATG |
| HC10 | TTGATCCAACTATGCGGG |

**Figure S1**

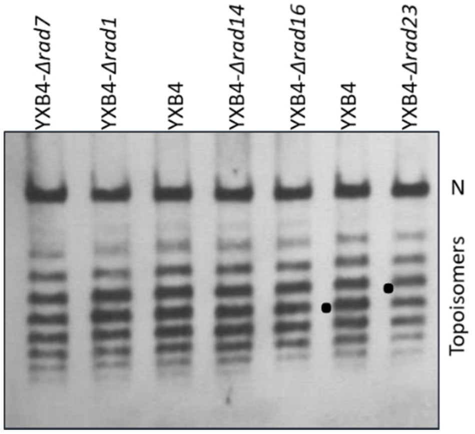


Figure S1. Analysis of topology of *HML* circles in *RAD* mutants. 40 µg of genomic DNA isolated from the indicated strains were separated by agarose gel electrophoresis in the presence of 30 µg/ml chloroquine and subjected to Southern blotting analysis. A representative experiment carried out twice is shown. *HML* topoisomers, nicked circles (N) and the Gaussian center of topoisomer distribution (dots) in WT (YXB4) and *rad23*Δ cells are indicated.
